# Tuber yield and processing quality response of French fried potatoes to nitrogen fertilization

**DOI:** 10.1101/759324

**Authors:** Lili Jiang, Ziquan Wang, Guanghui Jin, Guizhi Zhang, Chunyu Zhang

## Abstract

Nitrogen (N) is an important factor that influences potato production; appropriate N fertilizer management can optimize tuber yield and quality and thus reduce the risk of environmental N losses. The present study aimed to evaluate the effects of different nitrogen (N) application rates (0, 75, 150, 225, 300, and 375 kg ha^−1^) on the total tuber yield, marketable yield, dry matter content, reducing sugar content, sucrose content of tuber, and fry color index of two potato cultivars (Kennebec and Shepody) in 2016 and 2017. N supply significantly affected tuber yield and processing quality, but these effects depended on the year and/or cultivar. The results indicated that low (75 kg ha^−1^) and medium-N application rates (150 or 225 kg ha^−1^) had a positive effect on tuber yield and processing quality; however, high-N application rates (300 and 375 kg ha^−1^) resulted in lower yield and poor quality. Total and marketable yields responded quadratically to N, and they were optimized at 150 kg ha ^−1^ for Kennebec and 225 kg ha^−1^ for Shepody in both the years. The processing quality of tubers also responded quadratically with increasing N application rates. The optimal range of N application was approximately 145–185.83 kg ha^−1^ for Kennebec and approximately 93.44–288.67 kg ha^−1^ for Shepody according to the processing quality standards of French fried potatoes in China. To achieve the goals of high yield and high quality, N application rates should be 150 kg ha^−1^for Kennebec and 225 kg ha^−1^ for Shepody.

## Introduction

Potato (*Solanum tuberosum* L.) is a major global food crop because they can be grown under a wide range of environments and have high nutritional value [1]. In addition, potatoes can produce nutritious food more rapidly on less land and in harsh climates than other major crops. Most potatoes are grown for starch processing and French fry production, with a small proportion used for seed and table markets [2].

The processing of potatoes into French fries requires certain minimum quality attributes, including oblong to long tubers (> 75 mm), shallow eyes, minimal loss due to peeling, low reducing sugar, and light coloration [3]. Important factors influencing French fry grades and total tuber yield are nutrient management and choice of cultivar [4]. Kennebec and Shepody are widely grown in North China for French fries [5, 6].

Appropriate nitrogen (N) fertilizer management is a critical component for successful potato production [7, 8]. A sufficient N supply is important for achieving economically viable potato yields and the desired tuber size and quality targets from potato contracts [9, 10]. The availability of soil mineral N in planting can improve yields, but large doses of these soil minerals can delay tuber bulking, thereby leading to a decreased production of large tubers [11, 12]. For several cultivars, research conducted overseas has demonstrated that N nutrition can affect potato tuber quality [13–17].

Few studies have been conducted on the effects of N application on the processing quality of French fried potatoes. This research was initiated to investigate the effects of N management on potato yield and processing quality, including dry matter content, reducing sugar content, sucrose content of tuber, and fry color index. The specific objectives of the present study were to determine whether the range of N application rates applied to harvesting potatoes corresponded to specifications used for processing French fry products and to identify the optimal N application rate for Kennebec and Shepody cultivars. The results of the present study may help farmers planting potatoes for French fry processing and eventually improve the efficiency of N nutrition use and crop management.

## Materials and methods

### Ethics Statement

The experiments land is owned and managed by College of Agriculture, Heilongjiang Bayi Agricultural University, Heilongjiang Bayi Agricultural University permits and approvals obtained for the work and stu.The field studies did not involve endangered species or protected.

### Site description

The study was conducted in 2016 and 2017 at Yinlang Farm (124° 44’ 13.89" E, 56° 29’ 53.60" N) in the Daqing city of the Heilongjiang province, China. The soil at this site comprised sandy loam, and corn was previously grown at this site. Soil samples (at 0–20 cm depth) were collected before planting to determine their chemical characteristics (Table 1).

**Table 1.**
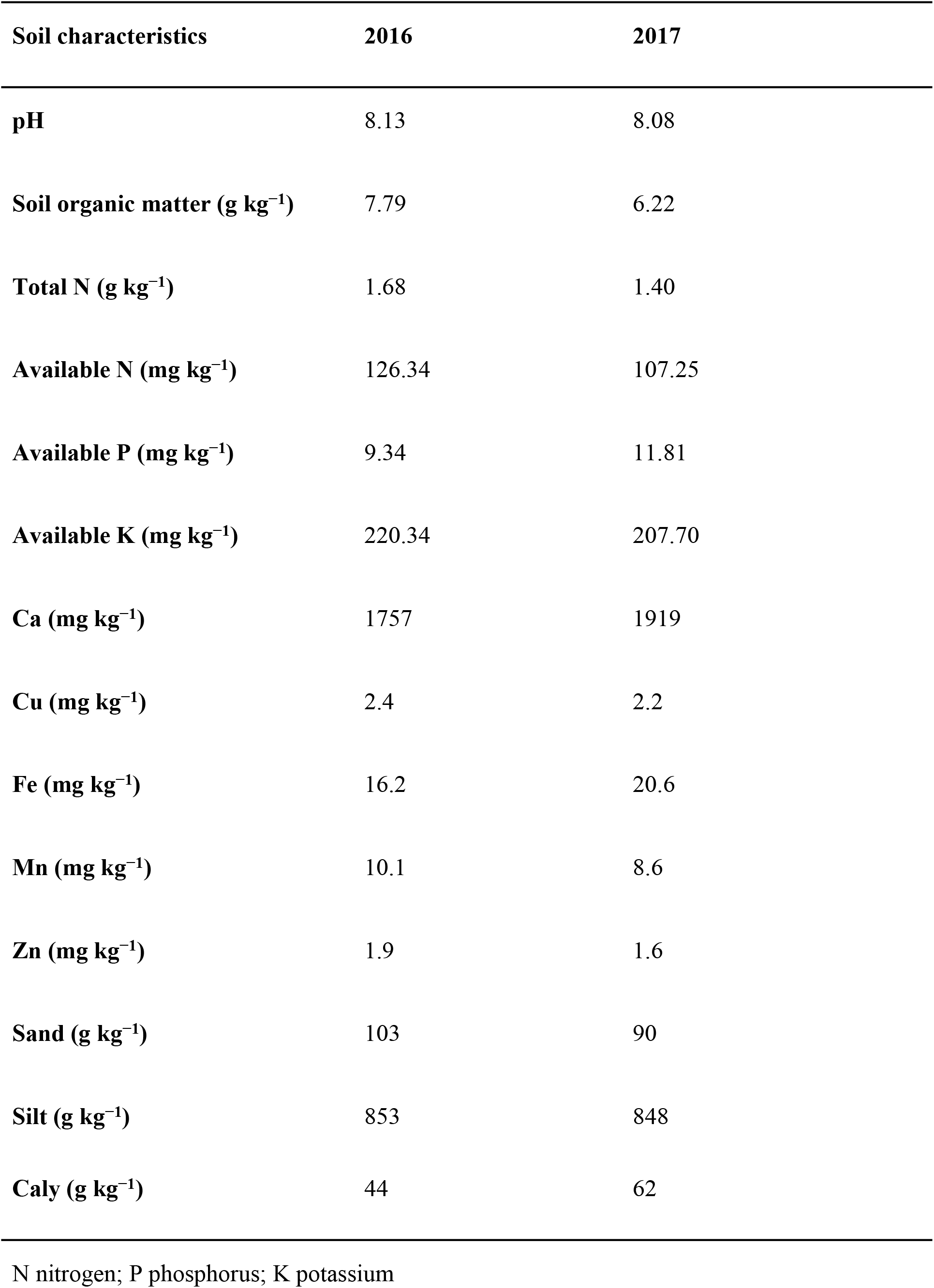
The chemical and textural characteristics of the soil samples at the experimental areas measured prior to the potato planting.

### Experimental design and treatments

All the experiments were conducted using a randomized complete block design with four replicates of each treatment. Treatments comprised Kennebec and Shepody cultivars and six N application rates (0, 75, 150, 225, 300, and 375 kg ha^−1^). The treatment with no N applied was considered a control, the 75 kg ha^−1^ was a low-N treatment, 150 and 225 kg ha^−1^ were medium-N treatment, and 300 and 375 kg ha^−1^ were high-N treatment. The plot size was 43.2 m^2^, with 8 m long rows and 0.90 m row spacing.

### Crop management

Potatoes were planted on May 5, 2016, and May 8, 2017, and the plant population was equivalent to 55555 plant^−1^. The source of N used was urea (46% N), and half of the N was applied as a base fertilizer and the remaining half was applied 45 days after introduction of each treatment. For all treatments, 180 kg P_2_O_5_ ha^−1^ (calcium phosphate) and 240 kg K_2_O ha^−1^ (K_2_SO_4_) were applied at planting.

Pest management and irrigation practices were conducted by research station personnel using common grower practices in the region. Rainfall and air temperature were measured using an automatic rain gage and thermometers installed at the experiment site. Water was applied when soil moisture reserves were reduced to 65% of the soil water-holding capacity. Water was applied at a rate of 6.8 mm h^−1^ with a portable overhead irrigation system.

### Tuber yield

The potatoes were harvested on September 25, 2016, and September 28, 2017. Sections (6 m long) from each of the four center rows of each plot were individually harvested, washed, and weighed, and tubers of 10 kg were selected from each plot for their quality assessment.

### Dry matter content

Dry matter content was calculated on the basis of specific gravity measurements. The specific gravity of marketable tubers was established by the weight in water, weight in air method on a composite of subsamples obtained from marketable tubers [18].

### Reducing sugar content

The concentration of reducing sugars in the potato tubers was determined using a colorimetric technique with dinitrosalicylic acid (DNS). Briefly, an extract was isolated from each potato tuber and was centrifuged to obtain pure juice; a 1:20 dilution (extract:distilled water) was then prepared with distilled water. The samples with the diluted extract were then supplemented with DNS at a ratio of 1:3 (diluted extract:DNS) and incubated in a boiling water bath for 5 min. The samples were cooled to room temperature, and their optical density was determined using a spectrophotometer at 540 nm as an excess of the optical density of the sample with the extract over the optical density of the background DNS solution. The obtained optical densities were recalculated to the concentrations of reducing sugars in g/L using calibration curves, and the dependences of the concentrations of reducing sugars on the irradiation dose were plotted for each stage [19].

### Fry color index

Five tubers were sampled from each replicate, and after removing 1–2 cm of each end, the tubers were peeled and five longitudinal strips (1 cm) were removed from the center of each tuber and fried for 3 min at 190°C. The average color of each of the 25 fries was visually rated immediately after frying according to the USDA frozen French fried potato color chart [20]. If the darker portions constituted more than approximately 50% of the strip, the strip would be given a higher rating. The chart ratings of 000, 00, 0, 1, 2, 3, and 4 represented the color of fried chips from light to dark, with 000 corresponding to the lightest color and 4 corresponding to the darkest color. The number of each grade of fried chips was counted as C000, C00, C0, C1, C2, C3, and C4. The equation of fry color index is as follows:

Fry color index = (C000 × 1 + C00 × 2 + C0 × 3 + C1 × 4 + C2 × 5 + C3 × 6 + C4 × 7) / 25

### Statistical analysis

Regression analysis was performed to determine the N application rate response curves for the measured potato traits (dependent variables). Significant regression equations with the highest coefficients of determination and that best explained the relationship between N application rates and dependent variables were selected and plotted using the software statistical package for the social sciences (SPSS) 16.0 package [21]. Every treatment was replicated four times. Data were subjected to analysis of variance and differences between means were tested at the 5% level using Duncan’s new multiple range test.

## Results

The total tuber yield was affected by cultivar, year, and N application rate. In addition, significant correlations were observed between cultivar and N application rate as well as between year and N application rate (Table 2). N application rate had a significant effect on the total tuber yield of Kennebec and Shepody cultivars in both 2016 and 2017. The total tuber yield of Kennebec and Shepody was lower in the absence of N fertilizer than in the presence of N fertilizer. N fertilizer application increased the total tuber yield in low- and medium-N treatments, but tuber yield decreased when N application rates were higher. The maximum total tuber yield was obtained with an N application rate of 150 kg ha^−1^ in 2016 and 2017 for Kennebec (41.35 and 48.47 Mg ha^−1^, respectively), whereas the maximum total tuber yield was obtained with an N application rate of 225 kg ha^−1^ for Shepody (38.46 and 41.08 Mg ha^−1^, respectively).

**Table 2.**
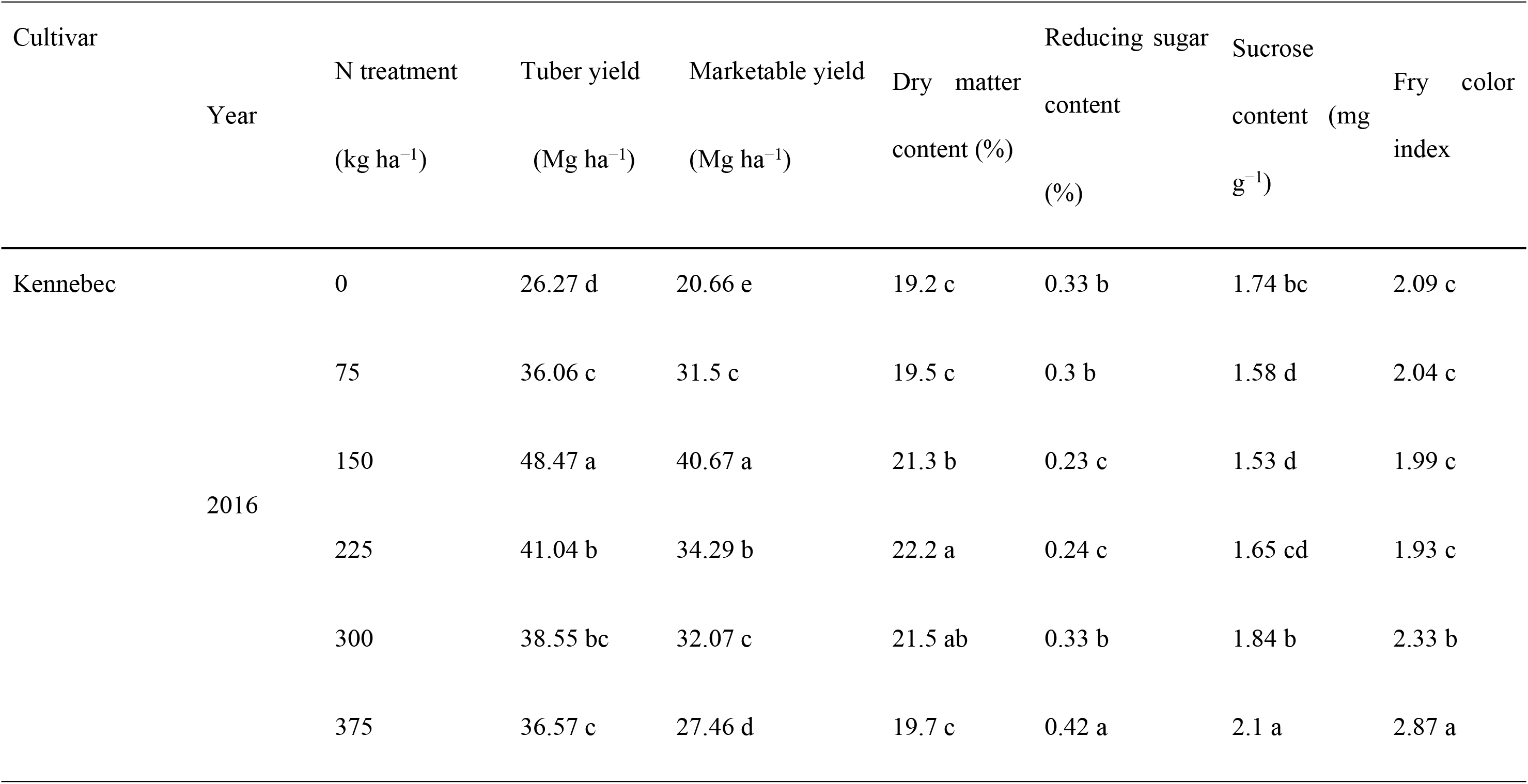
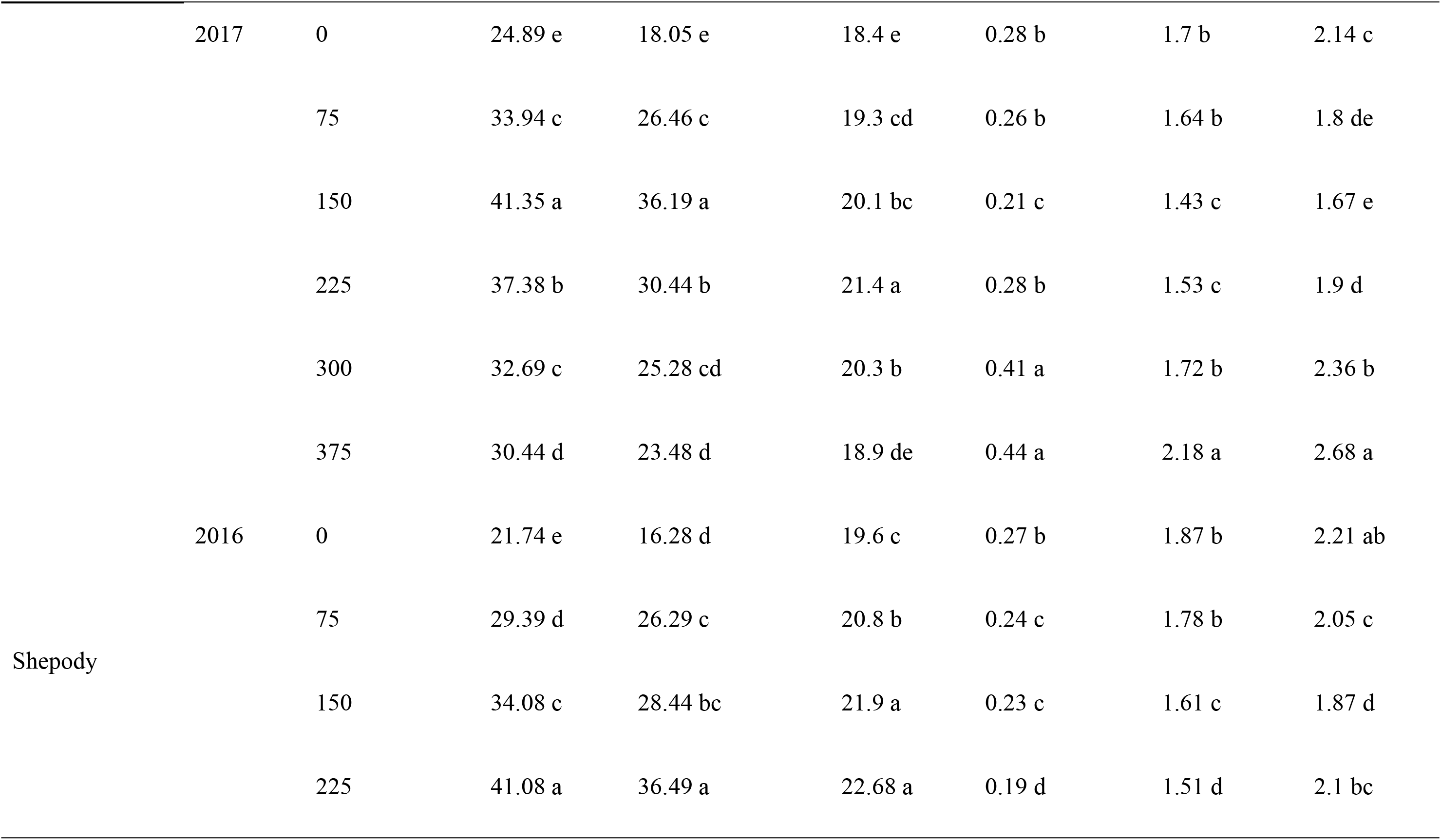
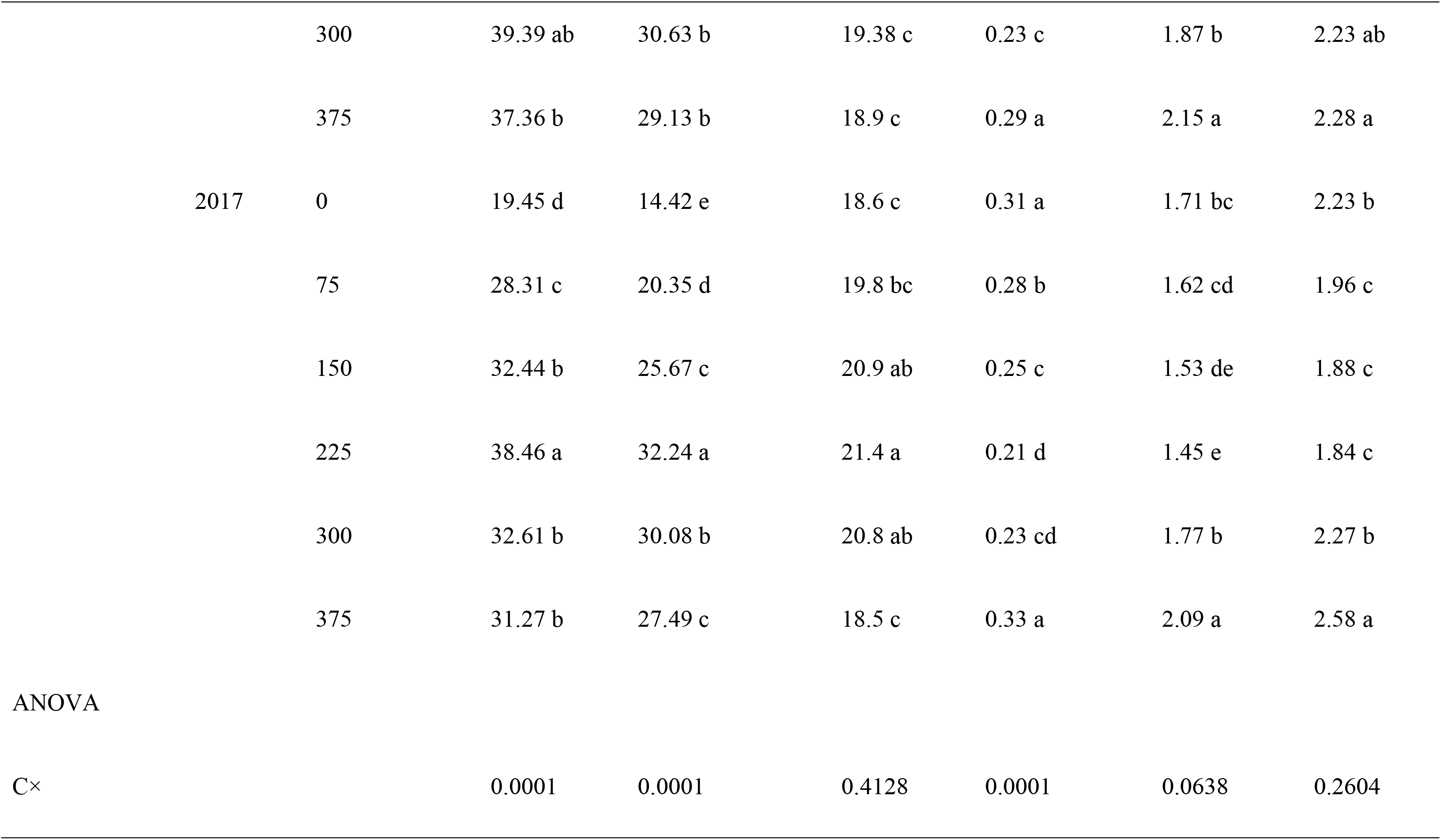
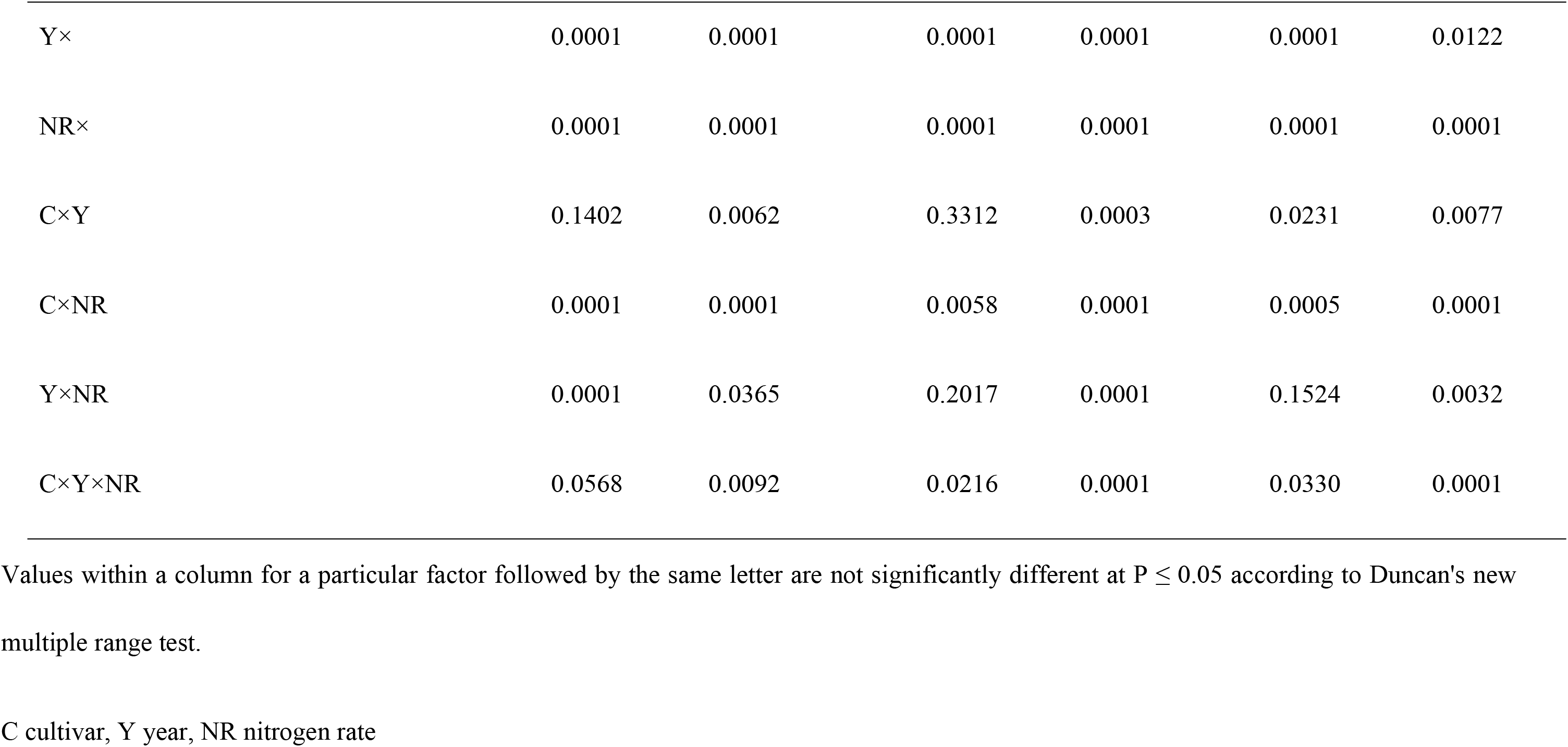
The effect of N application rates on tuber yield and processing quality of Kennebec and Shepody cultivars.

The marketable yield accounts for all tubers that can be sold for French fry production, after removing the small (< 7.5 cm), green, growth cracked, malformed, or decayed tubers. Cultivar, year, and N application rate, as well as cultivar and N interactions had no effects on marketable yield. The effects of N application on marketable yield were similar to those on total yield. Marketable yield responded quadratically with increasing N application rates in both Kennebec and Shepody cultivars across the two years (Table 2).

The dry matter content of the tubers was affected by year and N application rate. The dry matter content of Kennebec and Shepody increased in both the years when an N application rate of <225 kg ha^−1^ was applied (Table 2). High-N treatment had no positive effect on dry matter accumulation in tubers.

Reducing sugar content is a major quality factor in potato processing. In the present study, the reducing sugar content was affected by all the factors. However, the reducing sugar content of tubers did not differ between 0 and 75 kg ha^−1^ N treatments for Kennebec in 2016 or 2017, but low-N treatment affected the reducing sugar content of tubers of Shepody cultivar (Table 2). The N application rates of <150 kg ha^−1^ and 225 kg ha^−1^ decreased the reducing sugar contents in Kennebec and Shepody, respectively, both in 2016 and 2017.

The sucrose content of tubers was affected by year and N application rates. In 2016, an N application rate of 150 kg ha^−1^ resulted in the least sucrose content for Kennebec (1.43 mg g^−1^), with only slightly higher amounts in 2017 (1.53 mg g^−1^). There was no significant difference in the sucrose content in tubers between N application rates of 75 kg ha^−1^ in 2016 or 225 kg ha^−1^ in 2017. However, the N application arte of 150 kg ha^−1^ resulted in the lowest sucrose content for Shepody in both the years (1.43 and 1.53 mg g^−1^), which was significantly different from that for other N application rates (Table 2).

The fry color index was affected by N application rate, with a significant cultivar and N interaction (Table 2). A lower color index indicates better potato quality. An increase in the N application rate from 0 to 150 kg N ha^−1^ resulted in a decrease in the fry color index of Kennebec in 2016 and 2017, but an increase in the N application rate of >150 kg N ha^−1^ resulted in an increase in the fry color index (Table 1).

The relationship between N application rate and tuber yield and processing of Kennebec and Shepody can be expressed as a quadratic function (Fig. 1). By derivation of these equations, the optimal N application rate was obtained for quality traits (Table 3). Only potatoes that meet the quality index for standard French fry can be processed. The quality standards for French fry potatoes in China are as follows: dry matter content of tubers should be within 21%–23%, reducing sugar content of tubers should within 0.25%–0.35%, sucrose content of tubers should be within 1.50–1.7 mg g^−1^, and fry color index should ≤2.0 [22]. The range of N application rate for French fry potatoes was determined using the equation of each processing quality (Table 3). Based on the ranges of N application rates of quality traits, we can conclude that the optimal range of N application rate was 145–185.83 kg ha^−1^ for Kennebec and 93.44–288.67 kg ha^−1^ for Shepody. To achieve the goal of high yields and high quality, the N application rate should be 150 kg ha^−1^ for Kennebec and 225 kg ha^−1^for Shepody.

**Table 3.**
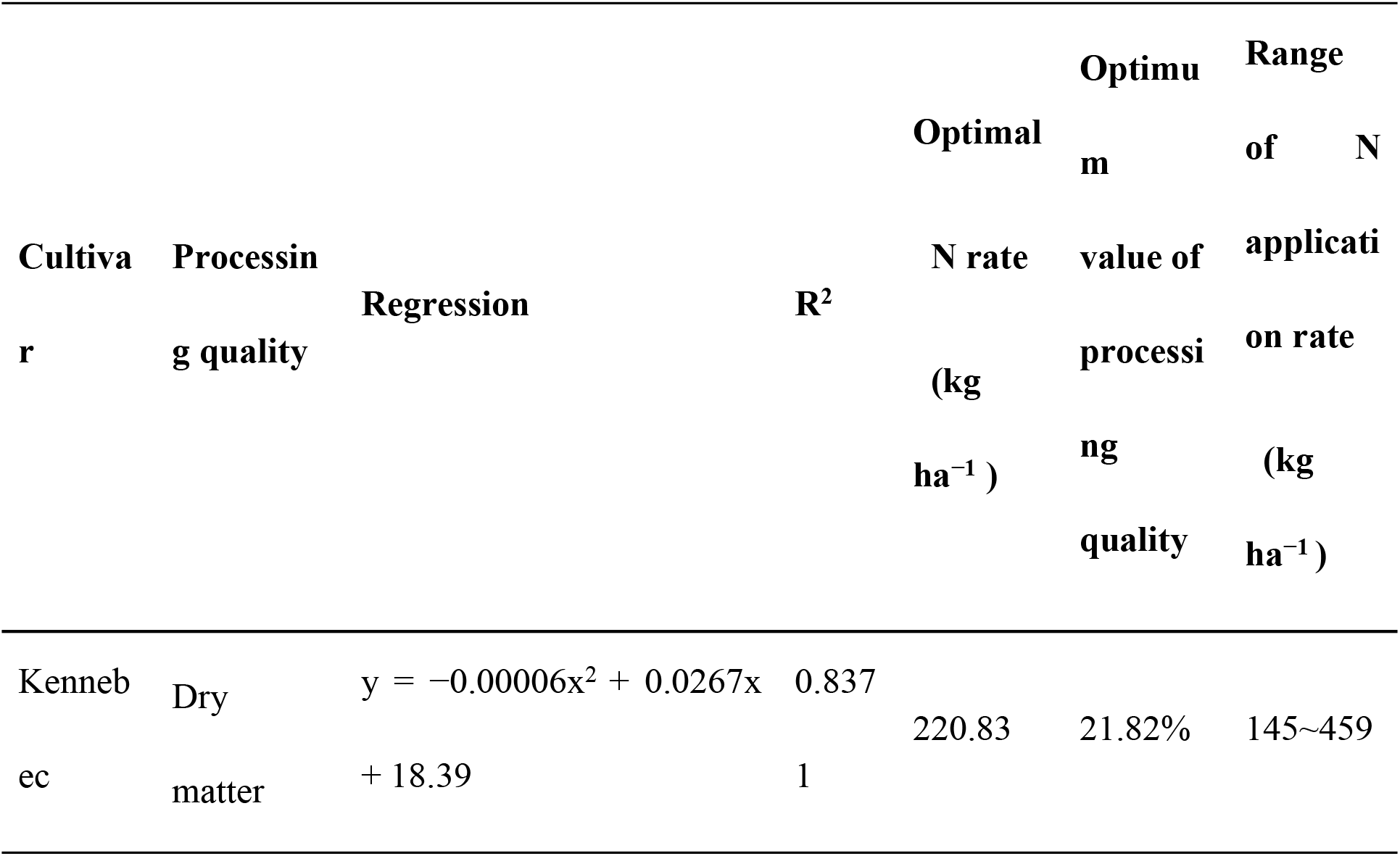
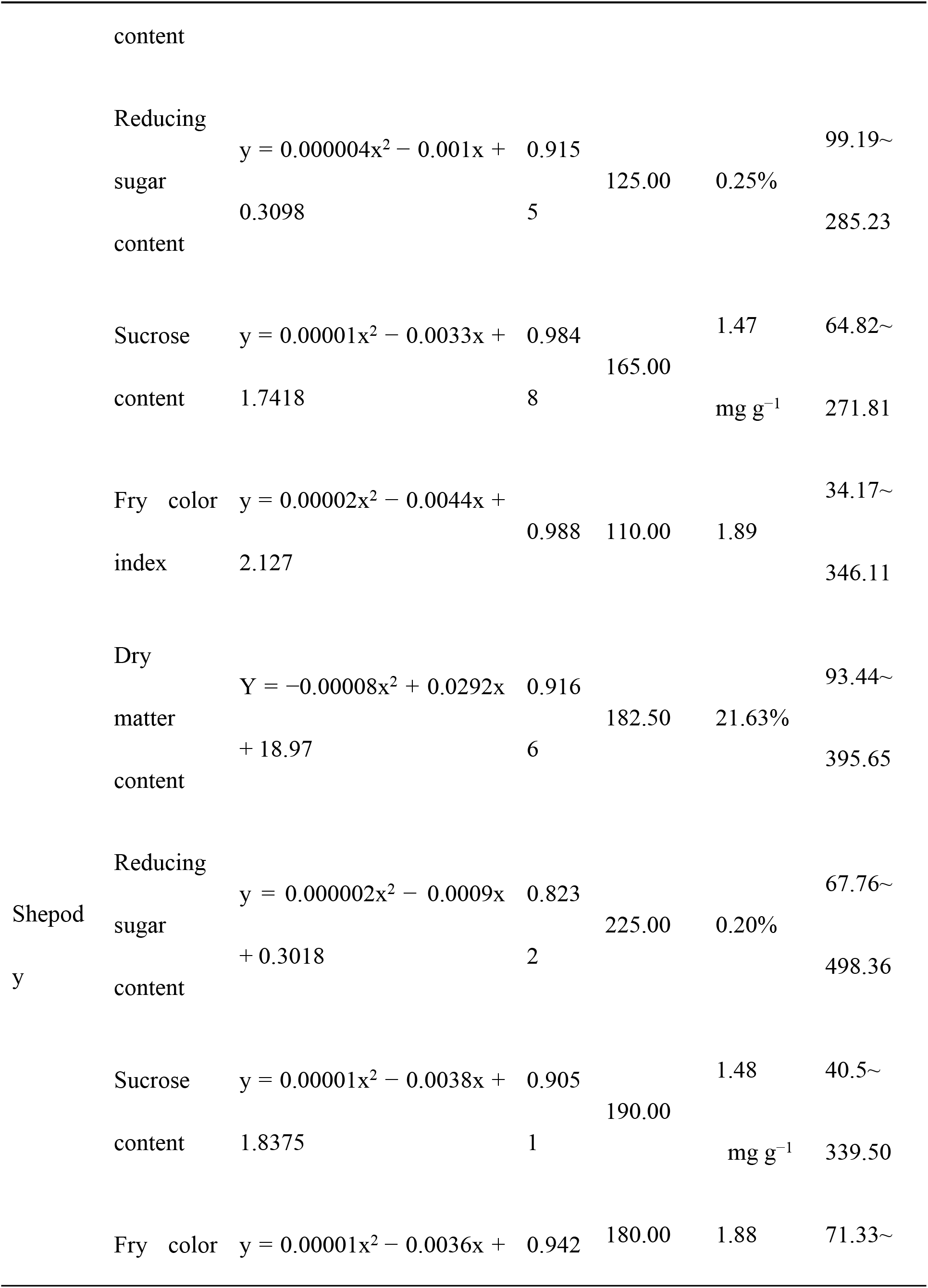
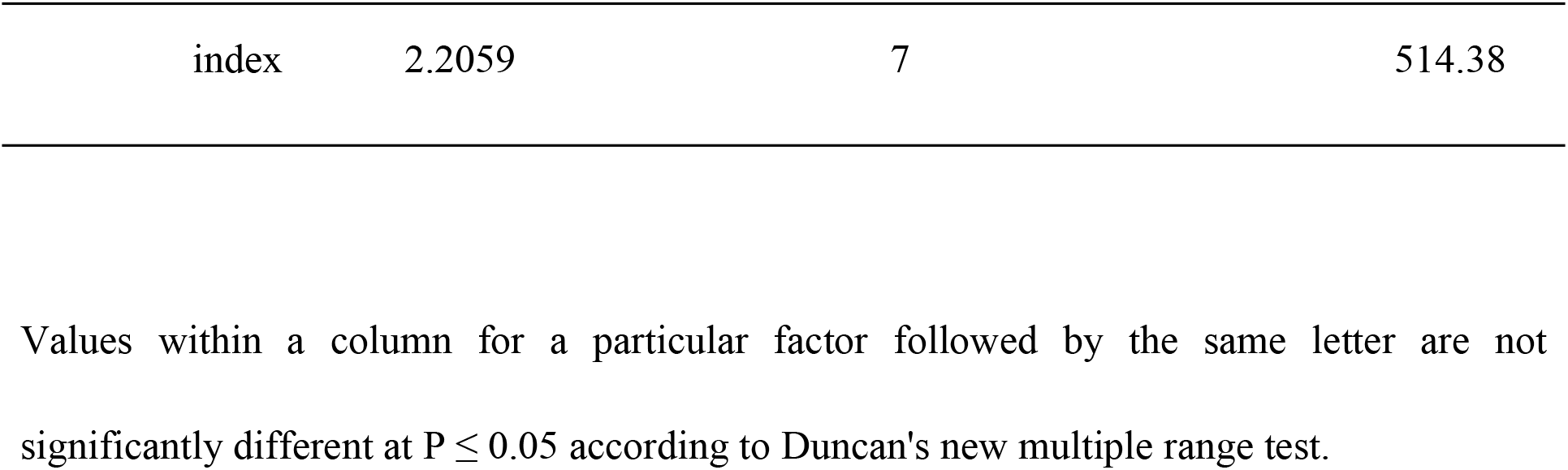
The optimal N application rate corresponded to the specification of processing for Kennebec and Shepody.

**Fig 1.**
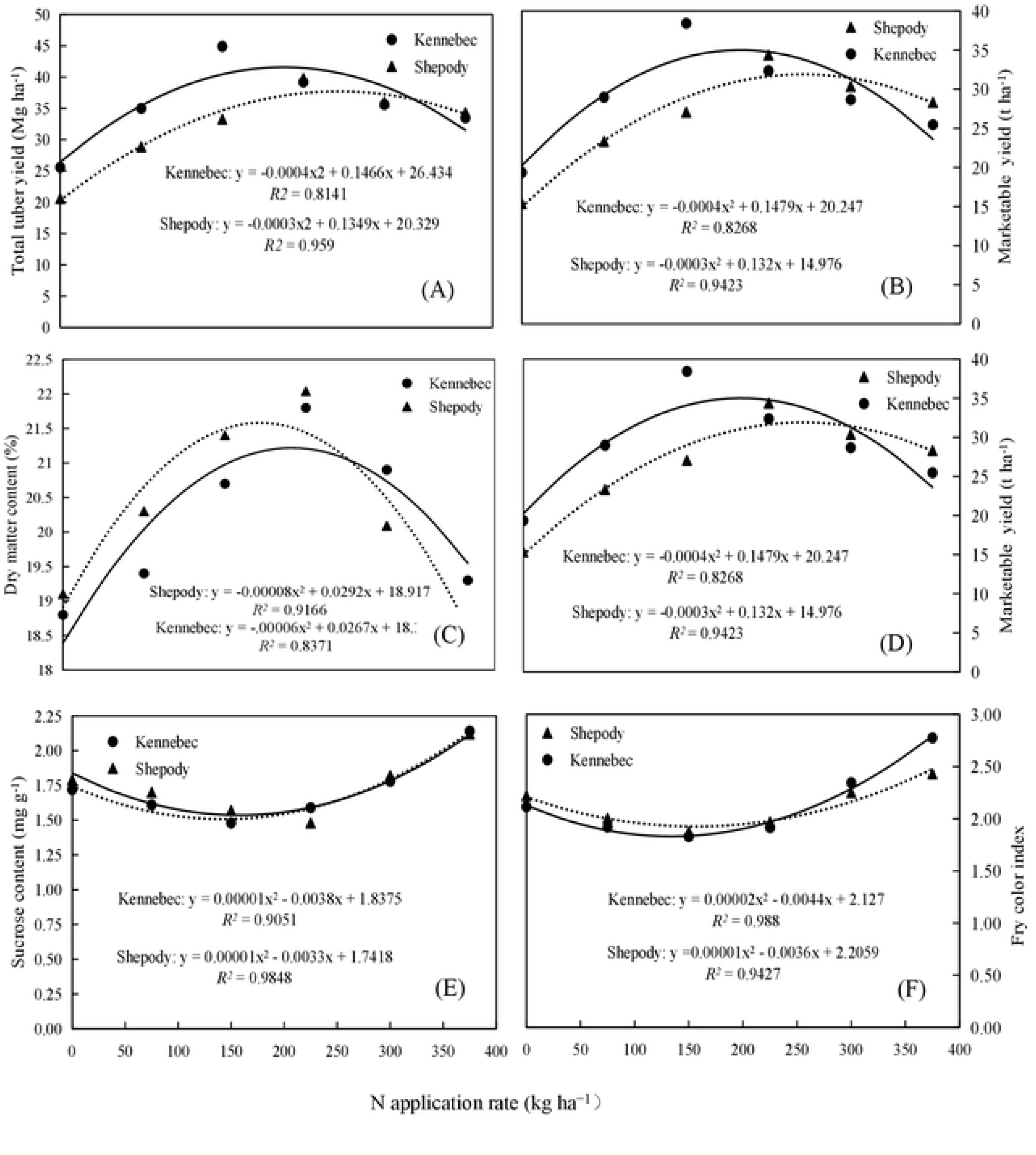
Relationship between N application rates and total tuber yields, marketable yield dry matter content, reducing sugar content, sucrose content of tuber, and fry color index of Kennebec and Shepody. The data presented are the average values of 2016 and 2017.

## Discussion

A previous study has reported that N supply significantly affected the total tuber yield and processing quality, but the amount of N applied was dependent on the year and/or cultivar [23]. A higher tuber yield and higher quality was achieved in the warm and longer growing season in 2016 than in the cooler, shorter growing season in 2017. The climate was more favorable for tuber bulking in 2016 than in 2017. Shepody demonstrated higher or comparable marketable yield and higher quality than the standard cultivar Kennebec. However, the tuber yield and quality traits were the best when the N application rates was 150 kg ha^−1^ for Kennebec and 225 kg ha^−1^ for Shepody. These results indicate that the efficiency of applied N use differs among cultivars. In addition to the specifics of each cultivar, marketable yield and quality must be considered for selection of French fried potatoes.

The effect of N on the growth and yield of potatoes has been the subject of several studies [24, 25], with an increase in potato tuber yield by N fertilization being a common finding [26–29]. Studies have reported that an increase in the N application rate increased the tuber yield, but the tuber yield did not always increase with an increasing N application rate. It has been demonstrated that excessive N reduces the tuber yield [30, 31]. The present study found that the maximum total tuber yield and marketable yield was obtained with the N application rates of 150 kg ha^−1^for Kennebec and 225 kg ha^−1^ for Shepody in both the years.

Dry matter accumulation in tubers is a result of the carbohydrates photosynthesized during tuber development that were retransmitted from leaves and stems [18]. The results of the present showed that the dry matter content increased in tubers when N was applied prior to the starch accumulation period; however, excessive N caused a decrease in the dry matter content and maturity of potatoes, which was in contrast with the findings of Sharifi et al. [32] who reported that N fertilization had no significant effect on tuber dry matter accumulation. Laboski and Kelling [33] have reported that tuber specific gravity decreases with increasing N application rates, particularly when the N rate exceeds the required N rate.

High reducing sugar content in the tuber causes darkening as a consequence of the mallard reaction and is therefore undesirable, and a low content of reducing sugar is preferred because it results in light and pleasing colors of desirable quality [34]. Dahlenburg et al. [5] have reported that the reducing sugar content in tubers was affected by the N application rate. In the present study, an increase in the reducing sugar content of Kennebec was observed in response to excessive N. The reducing sugar content of potatoes for French fry should range from 0.2% to 0.4% [22].

Sucrose is a good indicator of the chemical maturity of potatoes and is used to schedule harvests and monitor storage stress [6]. It has been demonstrated that increasing N fertilization can result in a decrease in the sucrose content at harvest, as well as in decreased effects on sucrose and increased effects on fructose accumulation in stored potatoes [35]. However, in this study, the sucrose content of Kennebec increased when the N fertilizer rate was higher than the optimal value.

Fry color index is an important indicator of food product characteristics because it is the essential quality parameter that is evaluated by consumers even before the food is consumed [36]. The N fertilizer rate has an inconsistent effect on chip or fry color [2]. Chip color has been reported to be unaffected by the N fertilizer rate [13, 16, 37, 38]. In the present study, N application did improve the fry color index when the N application rate increased from zero to the optimal N fertilizer rate (150 kg ha^−1^ for Kennebec and 225 kg ha^−1^ for Shpody); however, it resulted in a darker fry color with excessive N fertilization as reported in a previous study [25].

The range of N application rates was determined using the regression equation for dry matter, reducing sugar content, sucrose content of tuber, and fry color. The optimal range of N application was 145–271.81 kg ha^−1^ for Kennebec and 93.44–339.50 kg ha^−1^ for Shepody. It is remarkable that these ranges are basic standards for French fry potatoes. For example, if the dry matter content of a tuber is >21%, the tubers can also be used for French fries, but the dry matter content should not be <18%, indicating that the lower the reducing sugar content, sucrose content of tuber, and fry color index, the better is the quality of French fries.

## Conclusions

In the present study, we found that N supply significantly affected the tuber yield and processing quality of French fried potatoes, but these effects were dependent on the year and/or cultivar. Low- and medium-N application rates had a positive effect on tuber yield and processing quality; however, high N application rates resulted in lower yield and poor quality. The findings of our study demonstrated that the optimal range of N application was approximately 145–185.83 kg ha^−1^ for Kennebec and approximately 93.44–288.67 kg ha^−1^ for Shepody according to the processing quality standards of French fried potatoes in China. We suggest that the N application rates should be 150 kg ha^−1^ for Kennebec and 225 kg ha^−1^ for Shepody to obtain high yield and high quality.

## Acknowledgments

We thank the reviewers for their constructive suggestions for improving this manuscript; Hongyu Li and Xia Wang for the suggestion in the statistical analysis; and Feifei Xu for providing seed potatoes.

## Author Contributions

Conceptualization: ZiquanWang, Guanghui Jin

Formal Analysis: Lili Jiang, ZiquanWang, Guizhi Zhang

Funding Acquisition: Guanghui Jin

Investigation: Lili Jiang, Guizhi Zhang, Chunyu Zhang

Resources: ZiquanWang

Writing — Original Draft Preparation: Lili Jiang

Writing — Review & Editing: Lili Jiang, Ziquan Wang

